# Unraveling the genetic structure of the coconut scale insect pest (*Aspidiotus rigidus* Reyne) outbreak populations in the Philippines

**DOI:** 10.1101/726919

**Authors:** Joeselle M. Serrana, Naoto Ishitani, Thaddeus M. Carvajal, Billy Joel M. Almarinez, Alberto T. Barrion, Divina M. Amalin, Kozo Watanabe

## Abstract

The Philippines suffered from a devastating outbreak of the coconut scale insect pest, *Aspidiotus rigidus* Reyne inflicting significant economic losses to the country’s coconut industry. Despite the massive outbreak, little is known about the population and dispersal history of this invasive pest in the Philippines. Here, we examined the genetic diversity, structure and demographic history of *A. rigidus* sampled from localities with reported outbreaks from 2014 to 2017. We analyzed the genetic structure of seven *A. rigidus* outbreak populations using mitochondrial *COI* and nuclear *EF-1*α markers. Both markers and all methods of population genetic structure analyses indicate clear differentiation among the *A. rigidus* populations separating the northern (i.e., Luzon provinces) from the southern (i.e., Basilan and Zamboanga Peninsula) regions of the Philippines. Very low or no genetic differentiation was observed within and amongst the populations per geographic region indicating two unrelated outbreak events of the pest originating from two genetically uniform populations isolated in each respective region. Historical data supports the resurgence of an established *A. rigidus* population in the south which could have been driven by sudden climatic changes or human-induced habitat imbalance. Given no historical information, we disregard the possible resurgence from the northern population and infer that the outbreak could have resulted from a recent introduction of a non-native *A. rigidus* in the region. Our study provides valuable information on the genetic differentiation of the two *A. rigidus* groups that would be useful for developing and implementing biological control strategies against this pest in the Philippines.

## Introduction

Insect pest outbreaks are characterized by an explosive increase in the abundance of an insect population occurring over a relatively short period (Berryman, 1987). Large and rapid alterations in the environment or changes in the intrinsic genetic or physiological properties of individual organisms within a population can result to the resurgence of insect pests to outbreak-level status (Risch, 1987; Ziska *et al*., 2011). Likewise, insect outbreaks may occur when non-native species have no or few inefficient natural enemies, and if the local beneficial species are unable to suppress them in the area of introduction (Handley *et al*., 2011; Strayer *et al*., 2017). The invasive success of pest species may be determined by both the biology and environmental factors promoting its spread in a suitable area (Prentis *et al*., 2008; Renault *et al*., 2018). A better understanding of the source population, route and the mechanism of spread could provide valuable insights for designing and implementing quarantine strategies to understand the invasion success and decline of outbreak populations (Handley *et al*., 2011; Kobayashi *et al*., 2011).

In 2009, the Philippines suffered a devastating coconut scale insect (CSI) outbreak damaging the coconut palms on the provinces of Luzon (northern region of the Philippines) and currently on some areas in Mindanao (southern region) inflicting significant economic losses to the country’s coconut industry. The diaspidid insect *Aspidiotus rigidus* Reyne (Hemiptera: Diaspididae) resides on the underside of the leaf, blocks the stomata and sucks plant sap strongly reducing the plant’s photosynthetic activity leading to a characteristic yellowing and drying of the leaves. Severely infested coconut palms dry up and die within six months or less (Reyne, 1948). Prior to the renewed interest on *A. rigidus* due to the outbreak in the Philippines, historical and observational data on the spread of the invasive coconut pest has been scarce. Other than the most recent observation published by Watson *et al*. (2015), the last known study on the biology of this invasive species was conducted by Reyne (1947, 1948) with full documentation of the outbreak in the island of Sangi (North Celebes) in Indonesia from mid-1925 to 1928. The recorded past outbreak by Reyne (1948) naturally comes to an end after two years due to reduced female fecundity and high mortality of immature stages (Reyne, 1948). The decrease in *A. rigidus* population may have been associated with natural enemies that regulated the pest population overtime. However, the outbreaks from the localities infested with *A. rigidus* in the Philippines took longer times to recover (Watson *et al*., 2015), e.g., six years for the northern province of Batangas, or still on-going for the southern areas, i.e., Basilan, Zamboanga Peninsula, and the Caraga Region.

The introduction of *A. rigidus* to the Philippines, and its spread was believed to be either by wind or by accidental transportation of infested plants, coconut planting materials and products (Watson *et al*., 2015). Infestation in the northern provinces of the Philippines spread like wildfire from its initial local report in Tanauan, Batangas in the Calabarzon Region (Luzon) from 2009 reaching nearby coconut planted areas throughout the region. These outbreaks lasted for at least three years (Watson *et al*., 2015) and were reported manageable by 2015 (Manohar, 2015). The more recent outbreak in the southern region, specifically in Basilan, started early 2013 (Watson *et al*., 2015) implying its direct connection with the northern outbreak. However, given the means of spread by wind wherein crawlers are dispersed from one area to another (Watson *et al*., 2015), it is highly improbable for the infestation from the northern region to reach the infested southern islands moving pass other provinces planted with coconut palms along the way. Also, transport of infested plants from the northern provinces was highly unlikely given the national attention focused on quarantine and management strategies against the spread of the coconut scale insect during the outbreak (Javier, 2014; Manohar, 2015).

It is also likely that *A. rigidus* has been in the country as a minor pest and regulated by natural enemies. Based on historical reports, Lever (1969) reported sightings of *A. rigidus* in the Philippines, and Velasquez (1971) recounted that the pest was probably highly confined in the southern part of the country. It was more likely that the source of the sighted *A. rigidus* came from the island of Sangi in Indonesia given its relative closeness to Mindanao. In time, the immigrant *A. rigidus* could have established a resident population complemented by natural enemies limiting its colonization outside the area of introduction. Changes in anthropogenic, biotic interactions or climatic factors can influence a population’s rise to an outbreak level (Wilby & Thomas, 2002; Ziska *et al*., 2011). The recent outbreak observed in the southern part of the Philippines could have been caused by a sudden rise in the abundance of the supposedly established *A. rigidus* population due to factors such as human-induced habitat imbalance e.g., excessive use of pesticide affecting the natural enemies controlling the pest population, or climate change such as prolonged dry spell which may induce changes in the local biotic community.

Inference of the source population, route and the mechanism of spread of *A. rigidus* in the Philippines needs further assessment and confirmation. Tracing the history of an invasion or identifying the geographic origin of a pest population can be done by characterizing population-level genetic variation using molecular markers (e.g., Rugman-Jones *et al*., 2012; Kébé *et al*., 2016; Yang *et al*., 2017; Zhang *et al*., 2018). Sequencing selected gene fragments, e.g., mitochondrial COI is a traditional population genetic tool providing insights on dispersal pathways and population structure. The mitochondrial cytochrome oxidase (*mtCOI*) gene and the nuclear protein-encoding gene - elongation factor 1α (*EF-1*α) have been commonly used in studies investigating the origin (Provencher *et al*., 2005; Andersen *et al*., 2009), or inference of phylogenetic relationships (Andersen *et al*., 2010; Schneider *et al*., 2018) of various invasive diaspidid species.

Here, we aim to assess the population genetic structure and demography of the outbreak populations of the CSI, *A. rigidus* in the Philippines. Given the historical documentation of the pest in the southern region and the relatively extensive and rapid spread but faster recovery of the infestations in the northern region compared to the southern outbreaks i.e., Basilan and Zamboanga Peninsula, we hypothesize the presence of two distinct genetic groups for the outbreak events isolated within each geographic region. A population genetics approach is a useful tool to examine whether the northern and southern CSI outbreaks originated from immigrant or resident populations. To test the hypothesis, we utilized sequences of the mitochondrial cytochrome oxidase (*mtCOI*) gene and the nuclear protein-encoding gene - elongation factor 1α (*EF-1*α) to investigate the genetic structure and diversity of *A. rigidus* populations from localities with documented outbreak-level infestations in the Philippines from 2014 to 2017. Furthermore, we employed a coalescent genealogy approach to provide additional evidence on the demographic relationship of the outbreak *A. rigidus* populations between the northern and southern geographic regions in the Philippines.

## Materials and Methods

### Sample collection

*Aspidiotus rigidus* populations were sampled at seven localities with reported CSI outbreak across the Philippines from 2014 to 2017 (Fig. 1). The northern localities sampled were Orani, Bataan (BT; N14.769786, E120.454510), Nagcarlan (NG; N14.158930, E121.413670) and San Pablo (SP; N14.056420, E121.333300), Laguna, Tanauan (TN; N14.098870, E121.091330) and Talisay (TL; N14.093340, E121.010730) Batangas. The southern localities are Basilan (BS; N06.707853, E121.983358) and Zamboanga (ZB; N06.993166, E121.927963). See Table 1 for the more detailed information regarding location and sample collection information. Co-existence of other *Aspidiotus* species on the coconut palms sampled was possible, specifically *A. destructor* Sign. These two *Aspidiotus* species are difficult to separate morphologically, but some features of the live specimen and biology can be used to facilitate identification, i.e., the arrangement of eggs and egg skins relative to the insect’s body, scale cover appearance, and cuticle attributes (Reyne, 1948; Watson *et al*., 2015). Mature female scale insects were identified as *A. rigidus* based on the characteristic distribution of egg skins, which for this species occurs along the posterior or pygidial half of the insect body (Fig. 2). Non-parasitized adult females were carefully selected from infested leaves and preserved in 95% ethanol before the molecular analysis. To further confirm identification and the purity of samples, *A. rigidus* collected from Orani, Bataan were reared on *Garcinia mangostana* L. (mangosteen), a differential host of *A. rigidus* observed not to support multiple generations of *A. destructor* in the rearing facility of the Biological Control Research Unit (BCRU) located at De La Salle University (DLSU), Science and Technology Complex, Binan City, Laguna. A phylogenetic analysis was employed (see the succeeding molecular analysis below) to confirm the identifications of the field-collected samples by comparing it to *A. destructor* and mangosteen-reared *A. rigidus* sequences.

**Table 1.** Sampling localities of the outbreak *Aspidiotus rigidus* Reyne populations. N, number of individuals with *mtCOI* and *EF-1*α sequences; H, haplotypes indicated in Fig. 3.

| Locality |  | Code | Collection Date | *mtCOI* |  | *EF-1a* |  |
| --- | --- | --- | --- | --- | --- | --- | --- |
|  |  |  |  | N | H | N | H |
| Northern Region | DLSU-STC, Laguna<sup>a</sup> | AR | July 2017 | 35 | H<sub>1</sub> | 8 | H<sub>1</sub> |
|  | Orani, Bataan | BT | September 2015 | 21 | H<sub>1</sub> | 0 | H<sub>1</sub> |
|  | Nagcarlan, Laguna | NG | January 2015 | 24 | H<sub>1</sub> | 11 | H<sub>1</sub> |
|  | San Pablo, Laguna | SP | December 2014 | 21 | H<sub>1</sub> | 8 | H<sub>1</sub> |
|  | Talisay, Batangas | TL | December 2014 | 35 | H<sub>1</sub> | 18 | H<sub>1</sub> |
|  | Tanauan, Batangas | TN | December 2014 | 29 | H<sub>1</sub> | 8 | H<sub>1</sub> |
| Southern Region | Isabela, Basilan | BS | November 2016 | 44 | H<sub>2</sub> | 16 | H<sub>2</sub>; H<sub>3</sub> |
|  | Zamboanga City, Zamboanga | ZB | April 2017 | 96 | H<sub>2</sub> | 6 | H<sub>4</sub> |

<sup>a</sup>*Aspidiotus rigidus* Reyne reared on *Garcinia mangostana* L. at the DLSU-STC BCRU rearing facility from samples collected on the outbreak population in Orani, Bataan.

**Fig. 1.**
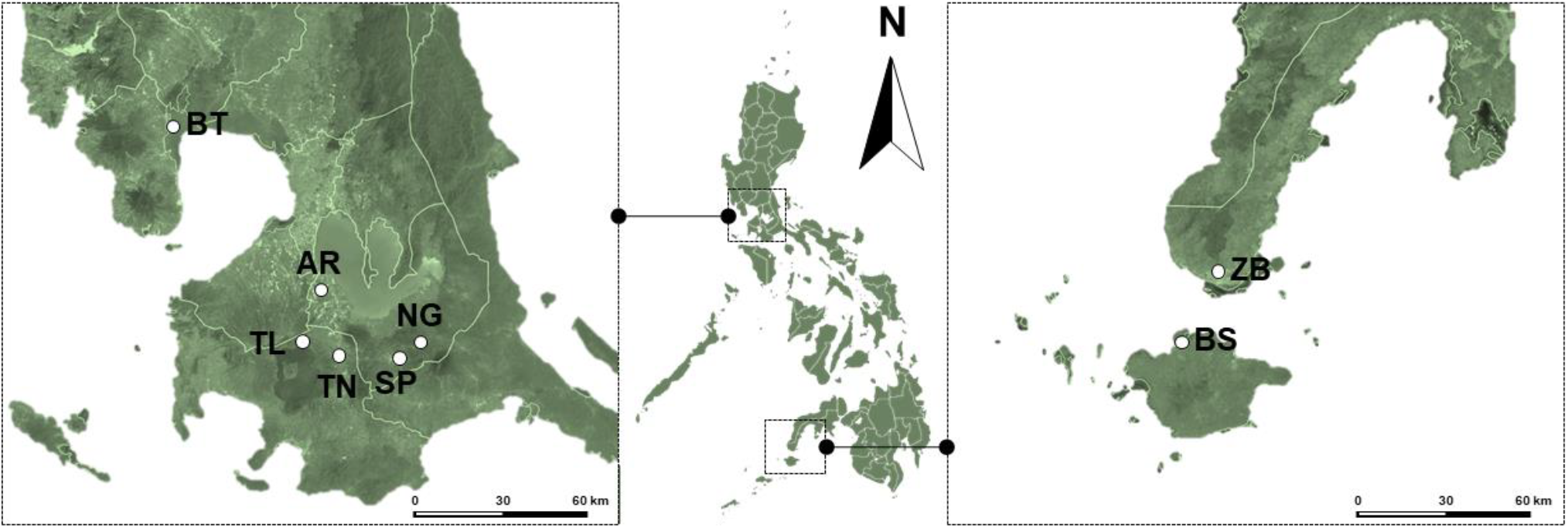
Map of the seven localities with reported *Aspidiotus rigidus* Reyne outbreak in the Philippines from 2014 to 2017. The insect rearing facility of the Biological Control Research Unit of De La Salle University labeled “AR”. Dots indicate sampling locations. Northern localities: Orani, Bataan (BT), Nagcarlan (NG) and San Pablo (SP), Laguna, Tanauan (TN) and Talisay (TL), Batangas; Southern localities: Basilan (BS) and Zamboanga (ZB). See Table 1 for the more detailed information regarding location and sample collection information.

**Fig. 2.**
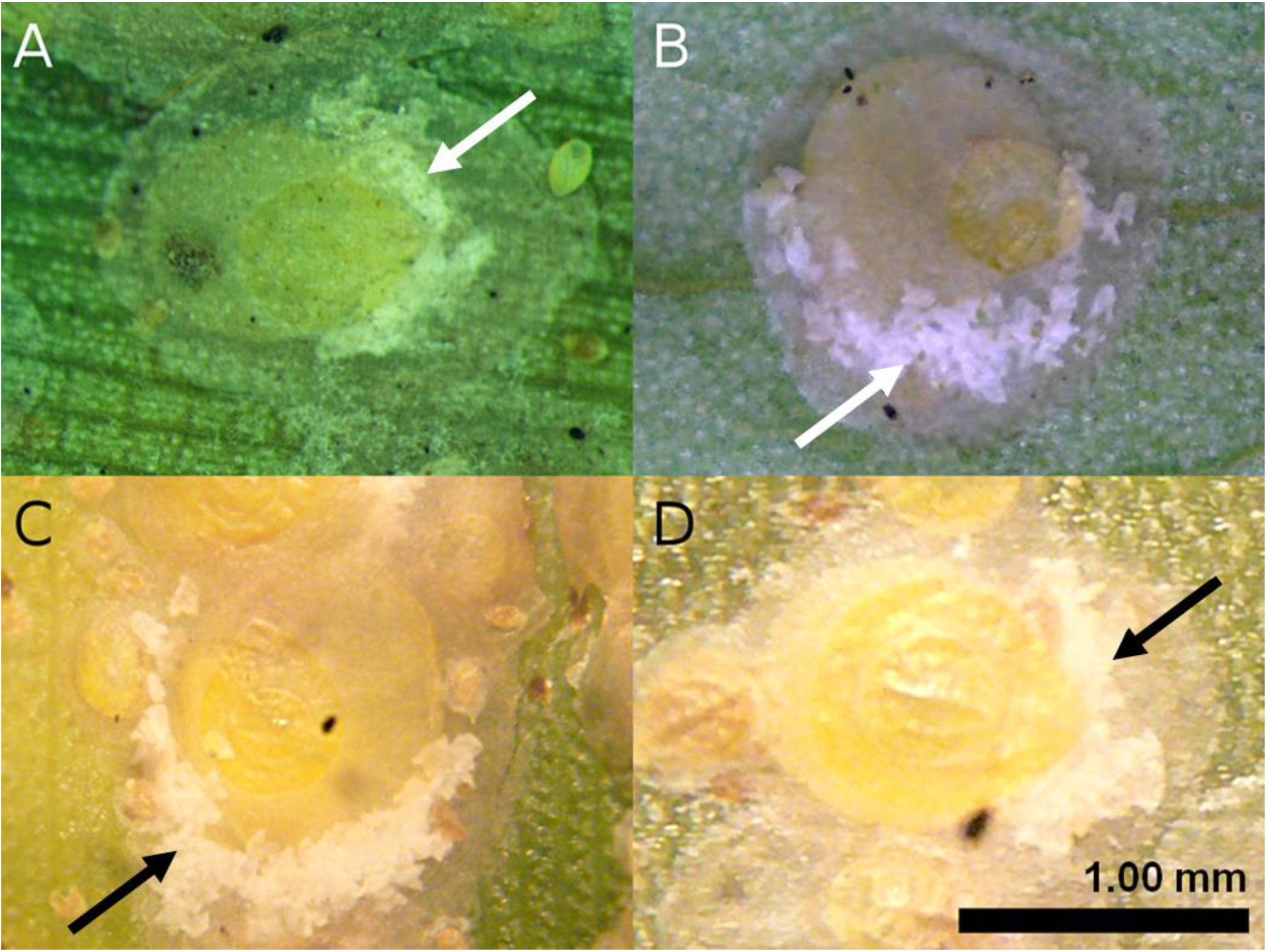
Representative adult female *Aspidiotus rigidus* Reyne from different outbreak areas: (A) Southern Tagalog Region (Laguna, Cavite, and Batangas); (B) Orani, Bataan; (C) Basilan; and (D) Zamboanga City. The arrows point to the egg skins, which for this species is characteristically distributed along the posterior or pygidial half of the insect body.

### DNA extraction, PCR amplification, and sequencing

Genomic DNA was extracted individually using the DNeasy Blood & Tissue Kit (QIAGEN, Hilden Germany) following the manufacturer’s guideline. Extraction was performed by crushing the insect body of each sample in individual microcentrifuge tubes using a micropestle. DNA concentration and quality were assessed by spectrophotometry (NanoDrop 2000 spectrophotometer, ThermoScientific).

The mitochondrial COI gene was amplified using the forward primer PcoF1 designed for scale insects by Park *et al*. (2010) and the standard reverse primer LepR1. The nuclear gene *EF-1*α was amplified using the forward primer *EF-1*α by Morse and Normark (2006) paired with the EF2 reverse primer (Palumbi, 1996). PCR reactions were performed in a 25 μl reaction containing 10× Buffer, 2.5 mM dNTP mixture, 25 mM MgCl_2_, 10 pmol of each primer, 1U of Taq DNA polymerase (TaKaRa Bio Inc.), and 2-50 ng of template DNA. PCR thermocycling was performed in a T100™ Thermal Cycler (Bio-Rad). Following the conditions from Park *et al*. (2011), the *mtCOI* gene was amplified with an initial denaturation step at 95°C for 5 min, followed by 5 cycles of 94°C for 40 s, annealing at 45°C for 40 s, extension at 72°C for 1 min and 10 s, and another 35 cycles of denaturation at 94°C for 40 s, annealing at 51°C for 40 s, extension at 72°C for 1 min and 10 s, and a 5 min final extension at 72°C after the last cycle. While, after an initial denaturation at 95°C, with a denaturation at 95°C for 30 s and extension at 72°C for 2 min every cycle, a touch-down procedure was performed for the amplification of the *EF-1*α gene following the protocol of Morse and Normark (2006) in which the initial annealing temperature of 58°C was decreased by 2°C every three cycles until a final temperature of 42°C was reached, then held for 18 cycles followed by a 5 min final extension at 72°C. PCR products were visualized in 1.5% agarose gels stained with Midori Green Direct (NIPPON Genetics Co. Ltd.), and cleaned using the QIAquick PCR Purification Kit (QIAGEN, Hilden, Germany). Samples were sent to Eurofins Genomics (Eurofins Genomics Co., Ltd.) for Sanger sequencing to produce both forward and reverse fragments.

### Genetic diversity and population structure

Sequences were assembled using CodonCode Aligner v. 5.1.5 (CodonCode Corporation). Before the subsequent molecular analysis, the sequences were aligned via MAFFT v. 7.409 (Katoh & Standley, 2013), and the ambiguously aligned regions were excluded using GBlocks 0.91b (Castresana, 2002). Sequence polymorphisms for both *mtCOI* and *EF-1*α gene were assessed. The number of variable sites (*S*) and haplotypes (*h*), average number of nucleotide difference (*k*), haplotype diversity (*Hd*), and nucleotide diversity (*Pi*) of the two marker genes were calculated in DnaSP v. 6.10.04 (Rozas *et al*., 2017).

The hierarchical analysis of molecular variance (AMOVA) was implemented in Arlequin v. 3.5.2.2 (Excoffier & Lischer, 2010). Two geographic groups were defined: The northern (Luzon) and the southern (Mindanao) groups prior to the analysis. Population pairwise *F*_*ST*_ was computed in Arlequin v. 3.5.2.2 using 1,000 permutations.

### Demographic inference

Tajima’s *D* and Fu’s *F*_*s*_ statistic tests were estimated to infer demographic history and dynamics in each population, for the two geographic groups, and for all populations grouped together in Arlequin v. 3.5.2.2 for both *mtCOI* and *EF-1*α datasets. Also, Fu and Li’s *D\** and *F\** test statistics were computed in DnaSP v. 5.0 to determine departures from the mutation-drift equilibrium (Fu & Li, 1993). Parameters of demographic expansion such as the moment estimators of time to the expansion *Tau*, effective population size before expansion (Theta0, *θ*_0_), effective population size after expansion (Theta1, *θ*_1_) between the observed and expected mismatches. The adjustment to a model of population expansion was estimated from the sum of squared deviation (*SSD*) and the raggedness index (*r*) in Arlequin v. 3.5.2.2.

### Gene flow analysis and median-joining networks of haplotypes

To test migration history between the two geographic groups, we calculated Bayes factors from the marginal likelihoods estimated in MIGRATE v. 4.4.0 (Beerli, 2005; Beerli & Palczewski, 2010) based on both *mtCOI* and *EF-1*α datasets. Migrate-n utilizes marginal likelihoods to compare and order structured population models (Beerli & Palczewski, 2010). The program provides estimates of historic gene-flow with the assumption that populations have reached mutation-migration-drift equilibrium. We tested eight possible models of migration history. Model 1 allows migration between the two groups, with the populations assumed to exist since a very long time. Model 2 presents a migration from northern to southern group, while model 3 presents vice versa with the populations assumed to exist since a very long time. Model 4 was one panmictic population encompassing the northern and southern groups. Model 5 allows divergence among populations within southern group splitting from the northern group, and migration from north to south, with the northern group existing for a long time and the southern group recently splitting off. Model 6 is a mirror image of model 5. Model 7 is similar to model 5 except that no interaction occurred between the two groups after the split. Model 8 is the vice versa of model 7. Similar parameters were used to run all models. A Bayesian search strategy was performed with the following parameters: one long chain (10,000 trees) with a burn-in of 5,000 iterations. A static heating scheme with 4 chains was applied using temperature parameters set by default with a swapping interval of one. Bayes factors were calculated via “*BF*” implemented in carlopacioni/mtraceR, a package for analyzing migrate-n outputs in R v. 3.5.2. Log Bayes factors of all models were calculated by comparing against the model that has the highest log-likelihood. The models are ranked based on LBF and calculated model probability.

Median-joining (MJ) networks (Bandelt, Forster, & Röhl, 1999) for the two markers were constructed to estimate the genealogical relationship in *A. rigidus* haplotypes via PopART v. 1.7 (Leigh & Bryant, 2015).

### Phylogenetic Analysis

Since no site variation was observed between the sequences in each population (except for the *EF-1*α sequences from Basilan), six representative samples per population, a total of 48 *mtCOI* sequences and 42 plus all 16 Basilan *EF-1*α sequences were chosen for the phylogenetic analysis. Sequences of *A. destructor* collected from coconut palms, identified based on the circular distribution of egg skins were selected as an outgroup (accession number: XX000000 for *mtCOI* and XX000000 for *EF-1*α). The Akaike information criterion corrected for sample size (AICc) was implemented to find the best fitting evolutionary model for phylogenetic reconstruction via jModelTest v. 2.1.10 (Darriba *et al*., 2012). The evolutionary model for the *mtCOI* sequences was TIM2+G, while TrNef was the model for the *EF-1*α sequences. Maximum likelihood (ML) tree inference was performed in RAxML-NG v. 0.5.1 (Kozlov at al., 2018) with 1,000 bootstrap replicates.

## Results

### Genetic diversity and population structure

All samples identified based on the characteristic distribution of egg skins were confirmed as *A. rigidus* by comparing the sequences with the *A. rigidus* reared on mangosteen and *A. destructor* sequences. DNA sequence analysis of all the concatenated 647-bp *mtCOI* sequences of 305 individuals from seven *A. rigidus* outbreak populations collected from 2014 to 2017 in the Philippines, with the mangosteen-reared samples revealed only two distinct haplotypes (*h*), separated by 31 polymorphic sites (*s*). Haplotype diversity (*Hd*) was calculated to be 0.050 +/−0.005 SD. Average number of nucleotide difference (*k*) was 15.447 and nucleotide diversity (*Pi*) was 0.024 +/−0.00025 SD. No sequence variation was found in the sequences of samples collected per populations. Reared *A. rigidus* and samples collected from the five populations of northern group, i.e., Orani, Bataan (BT), Nagcarlan (NG) and San Pablo (SP), Laguna, Tanauan (TN) and Talisay (TL), Batangas are grouped into one haplotype. Samples from the two populations of southern group, i.e., Basilan (BS) and Zamboanga (ZB) were grouped together in the second *mtCOI* haplotype. For the nuclear *EF-1*α gene, 75 concatenated sequences of 1007-bp length generated 14 polymorphic sites (*s*), with four haplotypes (*h*). Similar to the *mtCOI* sequences, all samples from the northern group clustered into one haplotype. For the southern group, samples from ZB grouped into one while BS were separated into two haplotypes. Haplotype diversity (*Hd*) was calculated to be 0.048 +/−0.064 SD. Average number of nucleotide difference (*k*) was 5.707 and nucleotide diversity (*Pi*) was 0.057 +/−0.00061 SD. Except for BS, all other localities have no sequence variation per locality.

Additionally, genetic diversity parameters have been calculated per geographic group. The *mtCOI* sequences for both groups, and the *EF-1*α sequence of the northern group showed no sequence variation. The *EF-1*α sequences from northern group had three haplotypes with an estimated *Hd* of 0.688 +/−0.039 SD, with a *k* value of 0.087 and a low *Pi* value of 0.00086 +/−0.00010 (Table 2). Both the median-joining haplotype network and ML inferred trees present distinct two and four haplotypes for the *mtCOI* and *EF-1*α dataset, respectively (Fig. 3 and S1).

**Table 2.** Parameters of genetic diversity and demographic analysis of the two population groups.

| Gene | Group | N<sub>1</sub> | S | h | Haplotype<sup>b</sup> | k | Hd (SD) | Pi (SD) | Tajima's D<sup>a</sup> | Fu's Fs | Fu and Li's D<sup>a</sup> | Fu and Li's F<sup>*</sup> |
| --- | --- | --- | --- | --- | --- | --- | --- | --- | --- | --- | --- | --- |
| *mtCOI* | Northern | 165 | — | — | H<sub>1</sub> | — | — | — | — | — | — | — |
|  | Southern | 140 | — | — | H<sub>2</sub> | — | — | — | — | — | — | — |
|  | **All** | **305** | **31** | **2** | — | **15.4465** | **0.4980 (0.005)** | **0.0239 (0.00025)** | **5.8452\*\*\*** | **59.5900** | **2.0420\*\*** | **4.4298** |
| *EF-1α* | Northern | 53 | — | — | H<sub>1</sub> | — | — | — | — | — | — | — |
|  | Southern | 22 | 2 | 3 | H<sub>2</sub>; H<sub>3</sub>; H<sub>4</sub> | 0.8701 | 0.6880 (0.039) | 0.0009 (0.00010) | 1.3276 | 0.9930 | 0.8506 | 1.1274 |
|  | **All** | **75** | **14** | **4** | — | **5.7067** | **0.4770 (0.064)** | **0.0057 (0.00061)** | **2.8256\*\*** | **12.8070** | **1.5502\*** | **2.3755** |

<sup>a</sup>Parameters with statistical test: \* indicates $p < 0.05$ ; \*\* indicates $p < 0.02$ ; \*\*\* indicates $p < 0.01$ .
<sup>b</sup>Haplotype data by DnaSP v. 6.10.04.

**Fig. 3.**
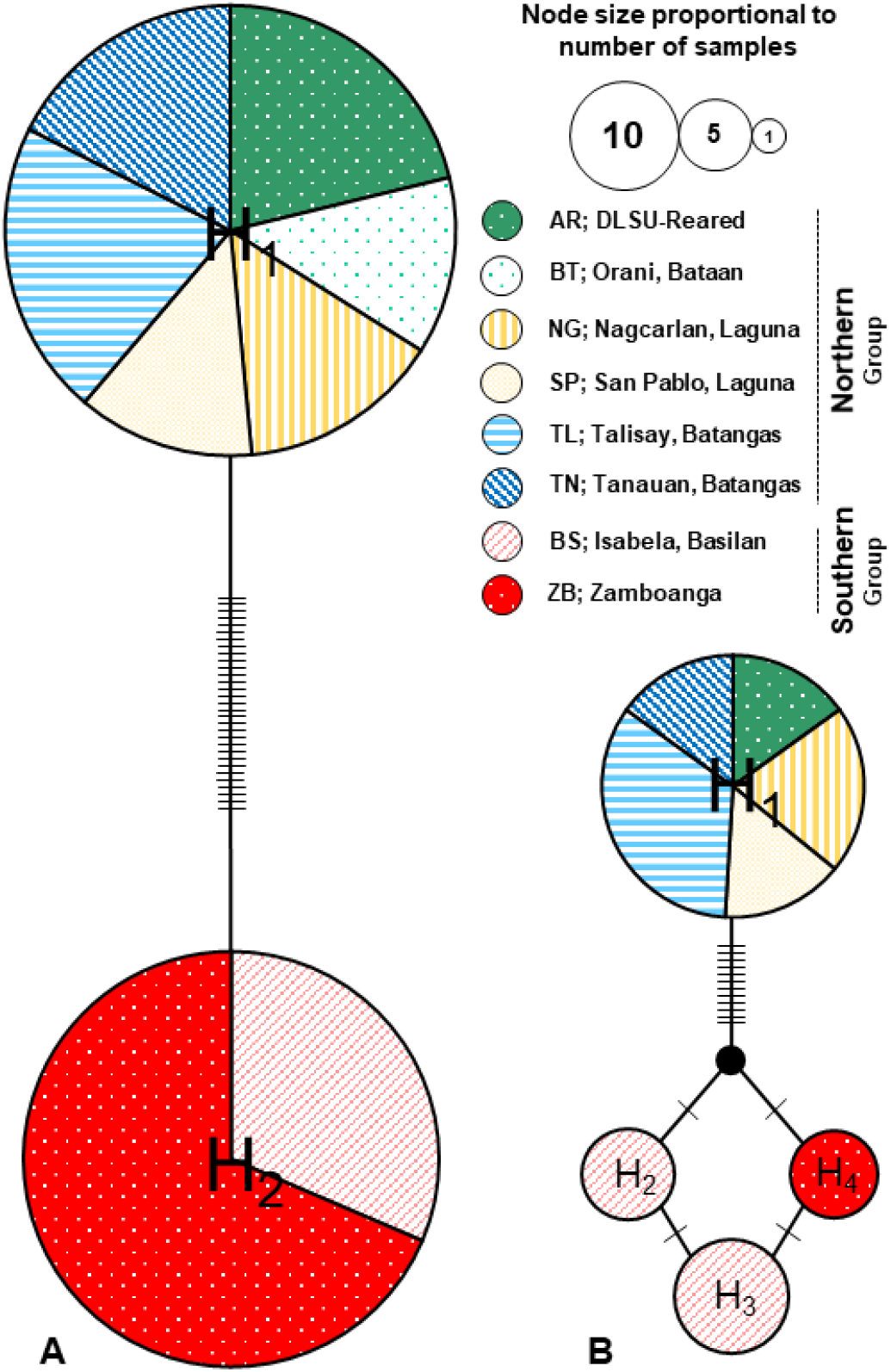
Median-joining network of the *Aspidiotus rigidus* Reyne populations from 305 individuals for the *mtCOI* gene (A), and 75 individuals for the protein-coding *EF-1*α gene (B), showing location and frequency of haplotypes. Each circle represents an observed haplotype; circle size indicates the number of individuals observed; the colors correspond to sampling localities. The total number of mutations, *Eta* presented as hatch marks.

AMOVA analysis indicated a highly structured genetic variability of 100% and 97.63% variations among the groups for *mtCOI* and *EF-1*α dataset, respectively. There were zero, or a relatively small percentage of variation among populations within groups and within populations. Except for the source of variation among populations within groups in the *mtCOI* data, AMOVA showed that significant genetic structure occurred in *A. rigidus* at various hierarchical levels (Table 3). Pairwise *F*_*ST*_ values varied from 0.00 to 1.00 for *mtCOI*, and 0.00 to 0.98 for the *EF-1*α dataset. The differentiation between populations was only significant when the comparison was between a northern and a southern population (Table S2). Moreover, pairwise *F*_*ST*_ values between the two groups showed a high value of 1.00 and 0.97 for the *mtCOI* and *EF-1*α dataset, respectively. As shown in the ML trees (Fig. S1), the phylogenetic analyses of both markers were consistent with the results of the analyses above. Samples clustered according to their geographic group, with the BS and ZB *EF-1*α sequences in three separate nodes.

**Table 3.** Partitioning of genetic variation at different hierarchical levels. * indicates p < 0.05; ** indicates p < 0.01.

| Gene | Source of Variation | d.f. | Sum of squares | Variance components | Percentage of variation | Fixation indices |
| --- | --- | --- | --- | --- | --- | --- |
| *mtCOI* | Among groups | 1 | 2347.869 | 15.50000 Va | 100 | F<sub>CT</sub>=1.00000* |
|  | Among populations within groups | 6 | 0 | 0.00000 Vb | 0 | F<sub>SC</sub>=0.00000 |
|  | Within populations | 297 | 0 | 0.00000 Vc | 0 | F<sub>ST</sub>=1.00000** |
| *EF-1α* | Among groups | 1 | 202.010 | 6.45405 Va | 97.63 | F<sub>CT</sub>=0.97630* |
|  | Among populations within groups | 5 | 5.199 | 0.09876 Vb | 1.49 | F<sub>SC</sub>=0.63040** |
|  | Within populations | 68 | 3.938 | 0.05790 Vc | 0.88 | F<sub>ST</sub>=0.99124** |

### Demographic history and gene flow

For both datasets, neutrality tests computation for all samples showed positive values for Tajima’s *D*, Fu and Li’s *D\**, Fu’s *Fs*, and Fu and Li’s *F\**, and were significant for the first two parameters (Table 2). Estimations per population for these parameters were mostly zero or positive but not significant, suggesting neither population expansion or purifying selection in these populations. Estimations of the *SSD* and *r* parameters both returns zero values, except for the *EF-1*α sequences from Basilan, with a significant *SSD* of 0.0283 (*p* < 0.001) and a not significant *r* of 0.2871. Other demographic parameters such as *Tau*, *θ*_0_ and *θ*_1_ index, are presented in Table S1. Results of the analysis in migrate-n were presented in Table S3. We found contrasting results for the two markers employed using Bayes factors to compare the eight models of dispersal. For the *mtCOI* dataset, model 7 was ranked best with a probability of 0.996. For the *EF-1*α dataset, model 3 was ranked best with a probability of 1.000.

## Discussion

We aim to describe the genetic structure and demography of the coconut scale insect pest *A. rigidus* from selected localities in the Philippines with reported heavy infestations collected from 2014 to 2017. Both the *mtCOI* and *EF-1*α markers and all methods of population structure analyses revealed strong differentiation among the *A. rigidus* populations separating the northern (Luzon) outbreak from the southern (Mindanao) region. The separation of the populations by geographic region and the observed lack of genetic variability within populations were represented graphically in the median-joining network and phylogenetic analysis employed in the study.

### Genetic structure of A. rigidus: Evidence of CSI “superclones” in the Philippines

Our results indicate the existence of two mitochondrial, and four nuclear haplotypes (one northern and three southern). Genetic population clusters result from multiple source populations contributing to an insect pest outbreak (Kobayashi *et al*., 2011). However, we only observed two clusters separating the outbreak populations into their respective geographic regions. Also, genetic variation was either absent or very low within and amongst the populations of the northern and the southern region, implying that populations from each region consisted of a single genotype. Hence, the presence of two distinct *A. rigidus* single genotype populations or “superclones” (Abbot, 2011) in the Philippines which supports our hypothesis on the occurrence of two genetically unrelated outbreak events in the country.

Several aspidiotine insects have obligate parthenogenetic populations (Normark & Johnson, 2011; Schneider *et al*., 2018). Accordingly, *A. rigidus* was observed to reproduce parthenogenetically. Yellow winged adult males are seen in outbreak populations but the sex ratio varies widely with males thought to be non-functional (Reyne, 1948; Watson *et al*., 2015). Parthenogenetic reproduction has been thought to be the leading driver to the dominance of “superclones” across space and time (Abbot, 2011). Similar to our findings, some invasive insect pests have been found to depend on clonal population structures to successfully invade and multiply in a broad range of niches. A highly specialized clonal genotype of a strictly asexual population of the pea aphid, *Acyrthosiphon pisum* Harris in central Chile was the main reason influencing the demographic success of the pest (Peccoud *et al*., 2008). Cifuentes, Chynoweth, and Bielza (2011) found no genetic variation and identified one single genetic type of the tomato leaf miner, *Tuta absoluta* Meyrick populations spreading through South America reaching the Mediterranean Basin. Likewise, a well-established invasive population of the oleander aphid, *Aphis nerii* B. de F. were reported having extremely low genetic diversity in the southern United States, with a “superclone” population supposedly obligatorily asexual (Harrison & Mondor, 2011). Caron, Ede and Sunnucks (2014) reported two widespread, invasive and strictly parthenogenetic “superclones” of the sawfly, *Nematus oligospilus* Forster dominating willows in three countries in the southern hemisphere i.e., South Africa, New Zealand, and Australia.

### Demographic history: Resurgence of resident population or recent introduction?

From historical reports, Lever (1969) claimed that the more invasive coconut scale insect *A. destructor rigidus* (now *A. rigidus*) reported by Reyne (1948) in Indonesia was also present in the Philippines. However, Velasquez (1971) did not disclose its occurrence across the archipelago and reported a highly probable confinement of the pest in the southern region of the country. Given this historical evidence, we assume that the southern populations have been existing for a long time. It could have supported our dispersal model for the mitochondrial sequences, except that the first reported sighting of *A. rigidus* in Tanauan, Batangas, Luzon was in 2009 (Watson *et al*., 2015) with no historical evidence of resident populations from the past. This suggests that the northern populations were most probably a recent introduction event from a different source.

A Bayesian search strategy was performed to assess the migration history between the northern and southern populations. However, our results from the mitochondrial and nuclear datasets are difficult to reconcile. We reiterate that in this historic gene-flow analysis, the two groups were assumed to have reached mutation-migration-drift equilibrium. Despite the contrasting results, both models indicate that sequences from the two groups do not belong to one panmictic population. However, given the difference in the divergence or migration pattern of the models for each marker, we were inconclusive in inferring the source of each outbreak population. Methodological assumptions (Knowles, Carstens & Keat, 2007) in the program Migrate-n, just like other coalescent-based approaches did not take into account another source of migrants, or that ancestral variation may come from populations that were not considered in the analysis. Hence, the inference of the possible source of the northern outbreak population needs further exploration.

On the other hand, lower genetic variation is expected for younger populations due to founder effects and genetic bottlenecks during colonization and establishment (Hewitt, 2004). Invasive or recently introduced species have been reported to exhibit reduced genetic variation (e.g., Tsutsui *et al*., 2000; Navia *et al*., 2005). Introduced populations are usually small so decreased genetic diversity is expected, and are often less variable than the source population which contributes to the invasive success of the species (Cifuentes, Chynoweth & Bielza, 2011). The nuclear marker revealed the existence of three southern haplotypes, with samples from Basilan having two distinct haplotypes. Genetic variation amongst the populations was very low and the Zamboanga *EF-1*α sequences are differentiated against Basilan with few nucleotide substitutions.

Genetic variation was already relatively low amongst the southern populations for the nuclear DNA, but in comparison to the uniformly genetic northern population, it indicates that the southern *A. rigidus* was relatively older in comparison to the northern region. Alongside previous historical reports, the level of genetic variation between the geographic regions supports our hypothesis of an existing resident *A. rigidus* population in the southern part of the Philippines. Local insect populations have the potential to outbreak due to anthropogenic and environmental changes (Berryman, 1987; Ziska *et al*., 2011). Similar observations on insect pests have been reported in literature. A notable example was by Kobayashi *et al*. (2011) which presented that the multiple nationwide outbreaks of the native populations of the mirid bug, *Stenotus rubrovittatus* Matsumura in Japan were induced by changes in the agro-ecosystem without invasion of populations from other areas. Populations of the pest were also genetically isolated by distance separated into genetic clusters occupying spatially segregated regions. Additionally, temporal fluctuations of pest insects in agroecosystems could be driven by various factors (Risch, 1987). Pesticide application may induce the resurgence of native pest insect populations by reducing the abundance of natural enemies or by the removal of competitive species in the area (e.g., Lu *et al*., 2010; Bommarco *et al*., 2011). Weather conditions can also trigger insect outbreaks due to the dramatic changes in pest abundance. Ward and Aukema (2019) reported that the cyclic outbreaks of the native tree-killing bark beetle, *Dendroctonus simplex* LeConte on tamarack in Minnesota, USA are climate-driven specifically associated with warmer and dryer years, more likely in areas with prior defoliation. Schwartzberg *et al*. (2014) simulated climate warming and observed warming-induced phenological shifts in the forest tent caterpillar, *Malacosoma disstria* Hübner about the phenology of its host trees. These findings illustrate the mechanisms by which anthropogenic and climatic changes induce outbreaks from native insect pests.

## Conclusion

The current opinion for the origin of the coconut scale insect outbreak in the Philippines was a recent introduction of *A. rigidus* from other countries of native range and spread via wind dispersal or importation of infested planting material from the northern region to the south given the timeline of the outbreak reports. However, our results indicate the separation of two distinct groups, the northern and southern *A. rigidus* from the outbreak populations collected from 2014 to 2017 in the Philippines. Very low or no genetic differentiation was observed within and amongst the populations per geographic region indicating two unrelated outbreak events of the pest species originating from two genetically uniform or “superclone” populations currently isolated in each respective region. Historical data supports our assumption on the current resurgence of an established *A. rigidus* population in the south. Given no historical information supporting the existence of an established *A. rigidus* populations in the northern region, we disregard the possible resurgence of a native population and suggest that the outbreak possibly resulted from a recent introduction of a non-native population. Assessment of the possible source population of the northern outbreaks needs further exploration.

The use of *mtCOI* and the nuclear *EF-1*α markers showed no or very low genetic differentiation for all *A. rigidus* populations. Other robust and more informative genetic markers such as microsatellites could provide further genetic information in studying the invasive coconut scale population. Further studies should also include more expansive sampling, taking into consideration other possible sources of *A. rigidus* such as Indonesia (Watson *et al*., 2015) and Vietnam (Schneider *et al*., 2018). This would provide a more robust and stringent population and gene flow estimation of *A. rigidus* in the Philippines. Nevertheless, our findings provided an initial important genetic basis and information for designing and implementing biological control strategies against the invasive CSI pest *A. rigidus* in the Philippines.

## Acknowledgements

We are grateful to Leslie Ann Ormenita and Reynaldo Majaducon of the Biological Control Research Unit of the Center for Natural Sciences and Environmental Research of De La Salle University for their technical support on the field work and preparation of samples. Many thanks to Dr. Maribet Gamboa for her valuable suggestions on the bioinformatics analyses conducted in this study. The authors acknowledge the support of the Sumitomo Electric Industries Group Corporate Social Responsibility Foundation, Ehime University’s Research Unit Program, and the Japan Society for the Promotion of Science (JSPS) Core-to-Core program (Asia-Africa Science Platforms).

## Author Contribution Statement

K.W., D.M.A., T.M.C., B.M.A. and J.M.S. conceptualized and designed the project; B.M.A., A.T.B., and D.M.A. conducted fieldwork; N.I. performed laboratory work; J.M.S. analyzed sequence data and drafted the article; A.T.B., D.M.A. and K.W. gave critical revisions; All authors approved the final version of the manuscript.

