## Supplementary Information for "Unraveling the genetic structure of the coconut scale insect pest (*Aspidiotus rigidus* Reyne) outbreak populations in the Philippines"

### Research Title

### Authors and Affiliations

Joeselle M. Serrana[1,3], Naoto Ishitani[1,3], Thaddeus M. Carvajal[1,3], Billy Joel M. Almarinez[2,3], Alberto T. Barrion[2,3], Divina M. Amalin[2,3] and Kozo Watanabe[1,3]

[1] Department of Civil and Environmental Engineering, Ehime University, Bunkyo-cho 3, Matsuyama, 790-8577, Japan
[2] Biology Department, College of Science, De La Salle University, 2401 Taft Avenue, Manila 1004, Philippines
[3] Biological Control Research Unit, Center for Natural Sciences and Environmental Research, De La Salle University, 2401 Taft Avenue, Manila 1004, Philippines

### Corresponding Author

Prof. Kozo Watanabe, PhD

Department of Civil and Environmental Engineering, Ehime University, Bunkyo-cho 3, Matsuyama, 790-8577, Japan
Phone & Fax Number: +81 (0) 89 927 9847

### ORCID ID

J. M. Serrana (0000-0002-6967-5407); T.M. Carvajal (0000-0001-5166-0058); B.J.M Almarinez (0000-0003-2562-9887); K. Watanabe (0000-0002-7062-595X)

#### Supplementary Figure

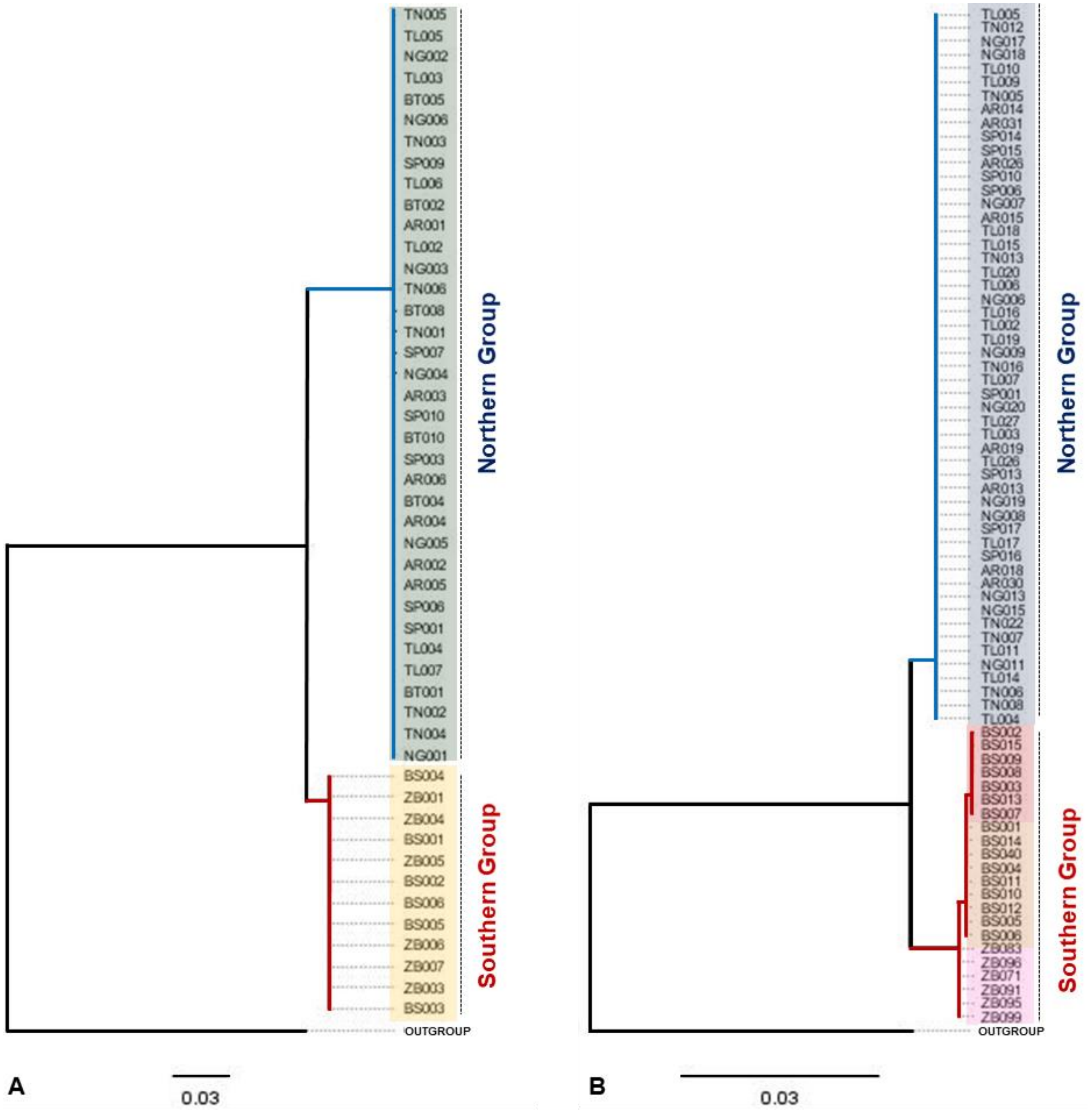


Fig. S1. Phylogeny of the outbreak *Aspidiotus rigidus* Reyne populations in the Philippines based on *mtCOI* (A) and *EF-1*$\alpha$ (B) sequences inferred using maximum likelihood. Refer to Table 1 for the corresponding code of each locality (e.g., BS003 sample collected from Basilan). Scale bar represents substitutions per site. Colors represent different haplotypes: *mtCOI* = 2; *EF-1*$\alpha$ = 4. Sequence of the morphologically similar coconut scale insect species, *Aspidiotus destructor* Signor*et al*so collected in the Philippines were included as outgroups.

#### Supplementary Tables

Table S1. Summary of the computations of the parameters for demographic expansion for the EF-1$\alpha$ dataset. The variance of the mismatch distribution is too small for the mtCOI dataset so no demographic parameters were estimated.

| Statistics | AR | NG | SP | TL | TN | BS | ZB | Mean | s.d. |
|---|---|---|---|---|---|---|---|---|---|
| Tau | 0.0000 | 0.0000 | 0.0000 | 0.0000 | 0.0000 | 0.7500 | 0.0000 | 0.1071 | 0.2835 |
| Tau qt 2.50% | 0.0000 | 0.0000 | 0.0000 | 0.0000 | 0.0000 | 0.0000 | 0.0000 | 0.0000 | 0.0000 |
| Tau qt 5% | 0.0000 | 0.0000 | 0.0000 | 0.0000 | 0.0000 | 0.0000 | 0.0000 | 0.0000 | 0.0000 |
| Tau qt 95% | 0.0000 | 0.0000 | 0.0000 | 0.0000 | 0.0000 | 0.0000 | 0.0000 | 0.0000 | 0.0000 |
| Tau qt 97.50% | 0.0000 | 0.0000 | 0.0000 | 0.0000 | 0.0000 | 0.0000 | 0.0000 | 0.0000 | 0.0000 |
| Thetaθ | 0.0000 | 0.0000 | 0.0000 | 0.0000 | 0.0000 | 0.0563 | 0.0000 | 0.0080 | 0.0213 |
| Thetaθ qt 2.50% | 0.0000 | 0.0000 | 0.0000 | 0.0000 | 0.0000 | 0.0000 | 0.0000 | 0.0000 | 0.0000 |
| Thetaθ qt 5% | 0.0000 | 0.0000 | 0.0000 | 0.0000 | 0.0000 | 0.0000 | 0.0000 | 0.0000 | 0.0000 |
| Thetaθ qt 95% | 0.0000 | 0.0000 | 0.0000 | 0.0000 | 0.0000 | 0.0000 | 0.0000 | 0.0000 | 0.0000 |
| Thetaθ qt 97.50% | 0.0000 | 0.0000 | 0.0000 | 0.0000 | 0.0000 | 0.0000 | 0.0000 | 0.0000 | 0.0000 |
| Theta1 | 0.0000 | 0.0000 | 0.0000 | 0.0000 | 0.0000 | 6833.4477 | 0.0000 | 976.2068 | 2582.8005 |
| Theta1 qt 2.50% | 0.0000 | 0.0000 | 0.0000 | 0.0000 | 0.0000 | 6823.4477 | 0.0000 | 974.7782 | 2579.0208 |
| Theta1 qt 5% | 0.0000 | 0.0000 | 0.0000 | 0.0000 | 0.0000 | 6823.4477 | 0.0000 | 974.7782 | 2579.0208 |
| Theta1 qt 95% | 0.0000 | 0.0000 | 0.0000 | 0.0000 | 0.0000 | 6823.4477 | 0.0000 | 974.7782 | 2579.0208 |
| Theta1 qt 97.50% | 0.0000 | 0.0000 | 0.0000 | 0.0000 | 0.0000 | 6823.4477 | 0.0000 | 974.7782 | 2579.0208 |
| SSD | 0.0000 | 0.0000 | 0.0000 | 0.0000 | 0.0000 | 0.0283 | 0.0000 | 0.0040 | 0.0107 |
| Model (SSD) p-value | 0.0000 | 0.0000 | 0.0000 | 0.0000 | 0.0000 | 0.0000 | 0.0000 | 0.0000 | 0.0000 |
| Raggedness index | 0.0000 | 0.0000 | 0.0000 | 0.0000 | 0.0000 | 0.2781 | 0.0000 | 0.0397 | 0.1051 |
| Raggedness p-value | 0.0000 | 0.0000 | 0.0000 | 0.0000 | 0.0000 | 1.0000 | 0.0000 | 0.1429 | 0.3780 |

Table S2. Population pairwise $F_{ST}$ values between the outbreak *Aspidiotus rigidus* Reyne populations for mtCOI and EF-1$\alpha$ data. Bold values are statistically significant at 0.05. Population code: AR, reared *A. rigidus* samples; BT, Bataan; NG, Nagcarlan; SP, San Pablo; TL, Talisay; TN, Tanauan; BS, Basilan; ZB, Zamboanga.

| mtCOI | AR | BT | NG | SP | TL | TN | BS | ZB |
|---|---|---|---|---|---|---|---|---|
| **AR** | | | | | | | | |
| **BT** | 0.0000 | | | | | | | |
| **NG** | 0.0000 | 0.0000 | | | | | | |
| **SP** | 0.0000 | 0.0000 | 0.0000 | | | | | |
| **TL** | 0.0000 | 0.0000 | 0.0000 | 0.0000 | | | | |
| **TN** | 0.0000 | 0.0000 | 0.0000 | 0.0000 | 0.0000 | | | |
| **BS** | **1.0000** | **1.0000** | **1.0000** | **1.0000** | **1.0000** | **1.0000** | | |
| **ZB** | **1.0000** | **1.0000** | **1.0000** | **1.0000** | **1.0000** | **1.0000** | 0.0000 | |

| EF-1a | AR | NG | SP | TL | TN | BS | ZB |
|---|---|---|---|---|---|---|---|
| **AR** | | | | | | | |
| **NG** | 0.0000 | | | | | | |
| **SP** | 0.0000 | 0.0000 | | | | | |
| **TL** | 0.0000 | 0.0000 | 0.0000 | | | | |
| **TN** | 0.0000 | 0.0000 | 0.0000 | 0.0000 | | | |
| **BS** | **0.9738** | **0.9769** | **0.9738** | **0.9818** | **0.9738** | | |
| **ZB** | **1.0000** | **1.0000** | **1.0000** | **1.0000** | **1.0000** | **0.7443** | |

Table S3. Bayes factors and log marginal likelihoods. The mod.rank indicates model ranking with “1” as the best model; mod.prob the value of probability.

| model | | Specification | lmL | LBF | mod.rank | mod.prob |
|---|---|---|---|---|---|---|
| mtCOI | 1 | xxxx | -1522.21 | -912.78 | 7 | 0.0000 |
| | 2 | x0xx | -1457.69 | -783.74 | 5 | 0.0000 |
| | 3 | xx0x | -1452.44 | -773.23 | 4 | 0.0000 |
| | 4 | x | -1809.00 | -1486.35 | 8 | 0.0000 |
| | 5 | x0Dx | -1472.43 | -813.22 | 6 | 0.0000 |
| | 6 | xD0x | -1433.60 | -735.55 | 3 | 0.0000 |
| | **7** | **x0dx** | **-1065.82** | **0.00** | **1** | **0.9960** |
| | 8 | xd0x | -1068.64 | -5.63 | 2 | 0.0040 |
| EF-1a | 1 | xxxx | -1595.40 | -38.70 | 5 | 0.0000 |
| | 2 | x0xx | -1586.16 | -20.22 | 2 | 0.0000 |
| | **3** | **xx0x** | **-1576.05** | **0.00** | **1** | **1.0000** |
| | 4 | x | -1596.00 | -39.90 | 6 | 0.0000 |
| | 5 | x0Dx | -1587.30 | -22.50 | 3 | 0.0000 |
| | 6 | xD0x | -1591.08 | -30.07 | 4 | 0.0000 |
| | 7 | x0dx | -1643.81 | -135.51 | 7 | 0.0000 |
| | 8 | xd0x | -1654.40 | -156.70 | 8 | 0.0000 |
